# The human α-defensin-derived peptide HD5(1-9) inhibits cellular attachment and entry of human cytomegalovirus

**DOI:** 10.1101/831594

**Authors:** Rebecca Böffert, Ramona Businger, Hannes Preiß, Dirk Ehmann, Vincent Truffault, Claudia Simon, Natalia Ruetalo, Klaus Hamprecht, Patrick Müller, Jan Wehkamp, Michael Schindler

## Abstract

Human cytomegalovirus (HCMV) infection causes severe illness in newborns and immunocompromised patients. Since treatment options are limited there is an unmet need for new therapeutic approaches. Defensins are cationic peptides, produced by various human tissues, which serve as antimicrobial effectors of the immune system. Furthermore, some defensins are proteolytically cleaved, resulting in the generation of smaller fragments with increased activity. Together, this led us to hypothesize that defensin-derived peptides are natural human inhibitors of virus infection with low toxicity. We screened several human defensin HNP4- and HD5-derived peptides and found HD5(1-9) to be antiviral without toxicity at high concentrations. HD5(1-9) inhibited HCMV cellular attachment and thereby entry and was active against primary as well as a multiresistant HCMV isolate. Moreover, cysteine and arginine residues were identified to mediate the antiviral activity of HD5(1-9). Altogether, defensin-derived peptides, in particular HD5(1-9), qualify as promising candidates for further development as a novel class of HCMV entry inhibitors.

**AUTHOR SUMMARY:** Defensins are peptides produced by various human organs which take part in the natural defense against pathogens. Recently, it has been shown that defensins are further cleaved to smaller peptides that have high intrinsic anti-microbial activity. We here challenged the hypothesis that these peptides might have antiviral activity, and due to their presumably natural occurrence, low toxicity. Indeed, we found one peptide fragment that turned out to block the attachment of the human cytomegalovirus (HCMV) to cells. Furthermore, this peptide did not show toxicity in various cellular assays or impede the embryonic development of zebrafish at the concentrations used to block HCMV. This is important, since HCMV is one of the most important viral congenital infections. Altogether, our results hold promise for the development of a new class of antivirals against HCMV.

## INTRODUCTION

Human cytomegalovirus (HCMV) is a β-herpesvirus with a high prevalence worldwide (1). Being non- or mildly pathogenic in immunocompetent individuals it causes severe disease in newborns, is the major pathogen of viral congenital infections and is a constant threat for immunocompromised patients, especially after organ transplantation (2). Established treatment strategies are based on the polymerase inhibitors Ganciclovir (GCV) and Foscarnet (PFA), but both frequently cause severe adverse-effects and may induce rapid drug-resistance (3). Additionally, the cytosine phosphonate inhibitor Cidofovir (CDV) used as second-line agent for drug resistant herpesvirus infections can induce multi-drug resistance leading to potentially lethal outcome (4). The introduction of the cytomegalovirus terminase inhibitor Letermovir inspired hope for novel alternative therapeutic regimens, which was dampened by the recent description of Letermovir resistance in multiple patients (3, 5). Altogether, there is a still unmet and urgent need for new approaches to treat human cytomegalovirus infection; especially in difficult to treat patients for instance organ and stem cell transplant recipients, pregnant women and their fetuses, or congenitally HCMV-infected newborns.

Defensins are anti-microbial peptides produced by all animal species (6, 7). They have a size of 16-50 aminoacids (aa), are amphipathic, rich in arginines and therefore have an overall positive charge. The secondary structure mainly consists of 3 β-strands that are stabilized by cysteine-build disulfide bonds. Depending on the size, abundance and overall structure they are categorized in α-, β- and θ-defensins. α-defensins are further subdivided in myeloid (HNP1-4) and enteric defensins (HD5 and HD6) (7, 8).

The antimicrobial activity of defensins is best characterized against bacteria (9). Defensins act in a multi-functional manner: they can penetrate membranes and form pores, and they interact with nucleic acids and glycosylated proteins (10–13). Furthermore it was shown that defensins are involved in the formation of anti-microbial NET-structures (14). While all these features contribute to the broad activity of defensins, they are restricted to certain defensin species and associated with the specific structural features and aa-motifs.

Although less studied, defensins also have antiviral activity (15, 16). For instance, α-defensins inhibit herpes simplex virus-2 (HSV-2) by interacting directly with viral particles or cellular heparan sulfates (17, 18). The α-defensins HNP1-3 interfere with HIV-1 glycoprotein binding to CD4, and the same defensins induce aggregation of Influenza-A virus and papillomaviruses (19–21). These examples show that α-defensins harbor antiviral activity against enveloped and non-enveloped viruses. However, the potential inhibition of HCMV by defensins has not been comprehensively studied thus far.

Recently it was shown that the enteric α-defensin HD5, but not HD6, undergoes proteolytic cleavage by human duodenal fluid resulting in the generation of HD5-derived peptides with potentially increased antimicrobial activity (22). We hypothesized that HD5-derived peptides might similarly have superior antiviral activity with low toxicity, as these molecules might naturally occur in humans. Indeed, here we identified HD5(1-9) as an attachment inhibitor of HCMV to various human cells with no toxicity at the concentrations in which they exert antiviral activity. Hence, defensin-derived peptides, and in particular HD5(1-9), are promising candidates for further development as a novel class of antiviral drugs.

## RESULTS

### Human α-defensins HNP4 and HD5 inhibit HCMV infection

We first aimed to assess the potential antiviral activity against HCMV of the α-defensins HNP4 (Fig. 1A) and HD5 (Fig. 1B) as well as defensin-derived peptides, occurring during natural proteolytic cleavage in human duodenal fluid (22) (Fig. 1). In a first approach, to assess potential toxicity, antiviral activity, as well as dose-dependency, we tested the defensins at two concentrations, high (75 µM) and low (7.5 µM) (Fig. 2). We infected primary human foreskin fibroblasts (HFF) with an HCMV-GFP reporter virus (MOI = 0.5) and added the peptides simultaneously with the infectious virus to the cells. At 40 hours post infection (hpi), cells were fixed, stained with DAPI, and the infection rate was calculated by automated cell counting (Fig. 2A, % GFP+/DAPI+ cells). Both full-length α-defensins, HNP4 and HD5, inhibited HCMV infection at 75 but not at 7.5 µM. Similar activity was observed for the short fragments HD5(1-9), HD5(7-32) and to a lesser extent for HNP4(1-11). Two peptides, HD5(1-9mod) and HNP4(1-11mod), were modified to protect them from proteases and increase activity by using D-aminoacids and by adding an acetate moiety to the N- and an amide moiety to the C-terminus (Fig. 1). These two defensin-fragments also blocked HCMV-infection at 75 µM (Fig. 2A).

**Figure 1:**
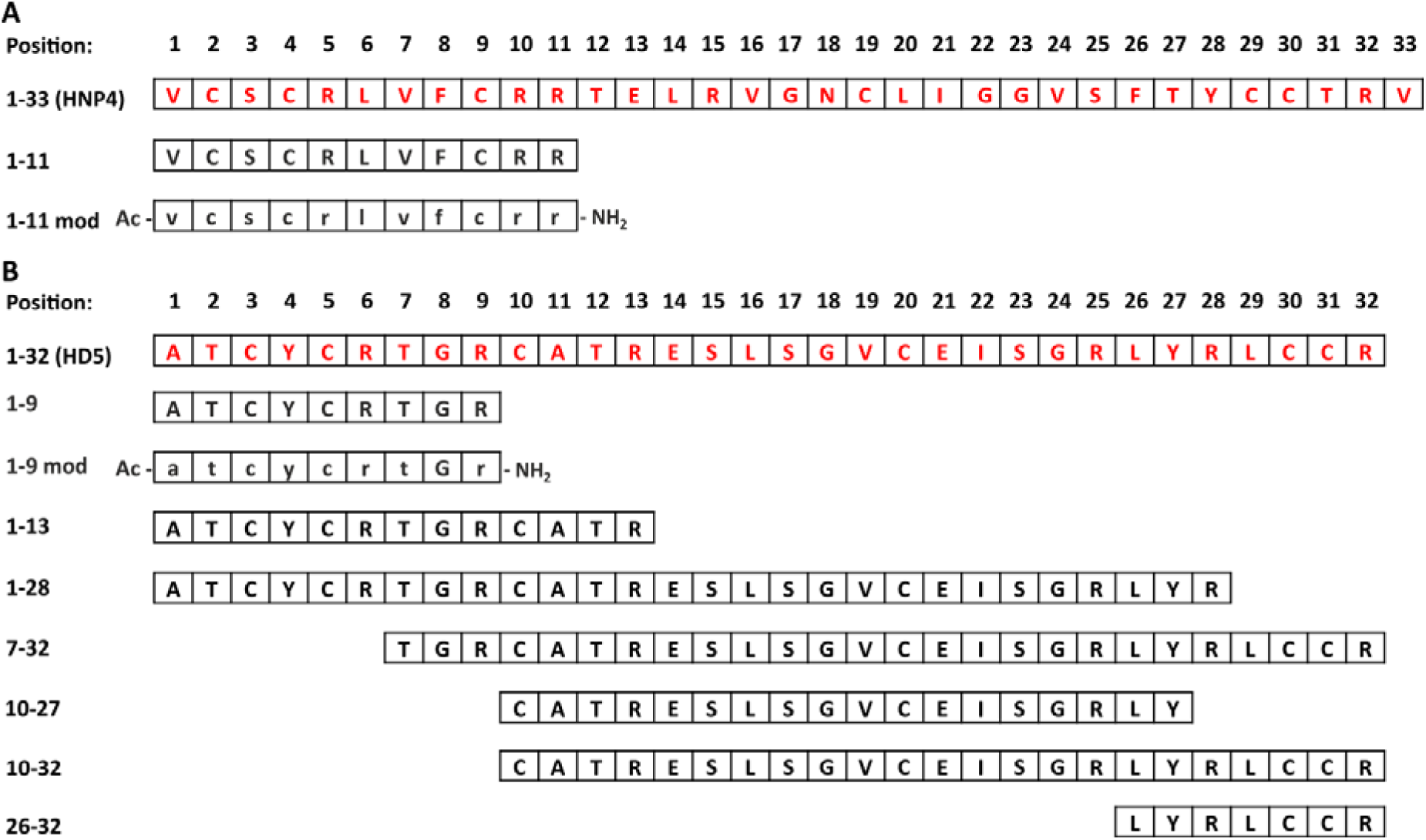
Primary amino acid sequence alignment of HNP4, HD5 and all corresponding fragments. (A) HNP4 or (B) HD5 full length proteins are highlighted in red. D-amino acids appear in lower case letters.

**Figure 2:**
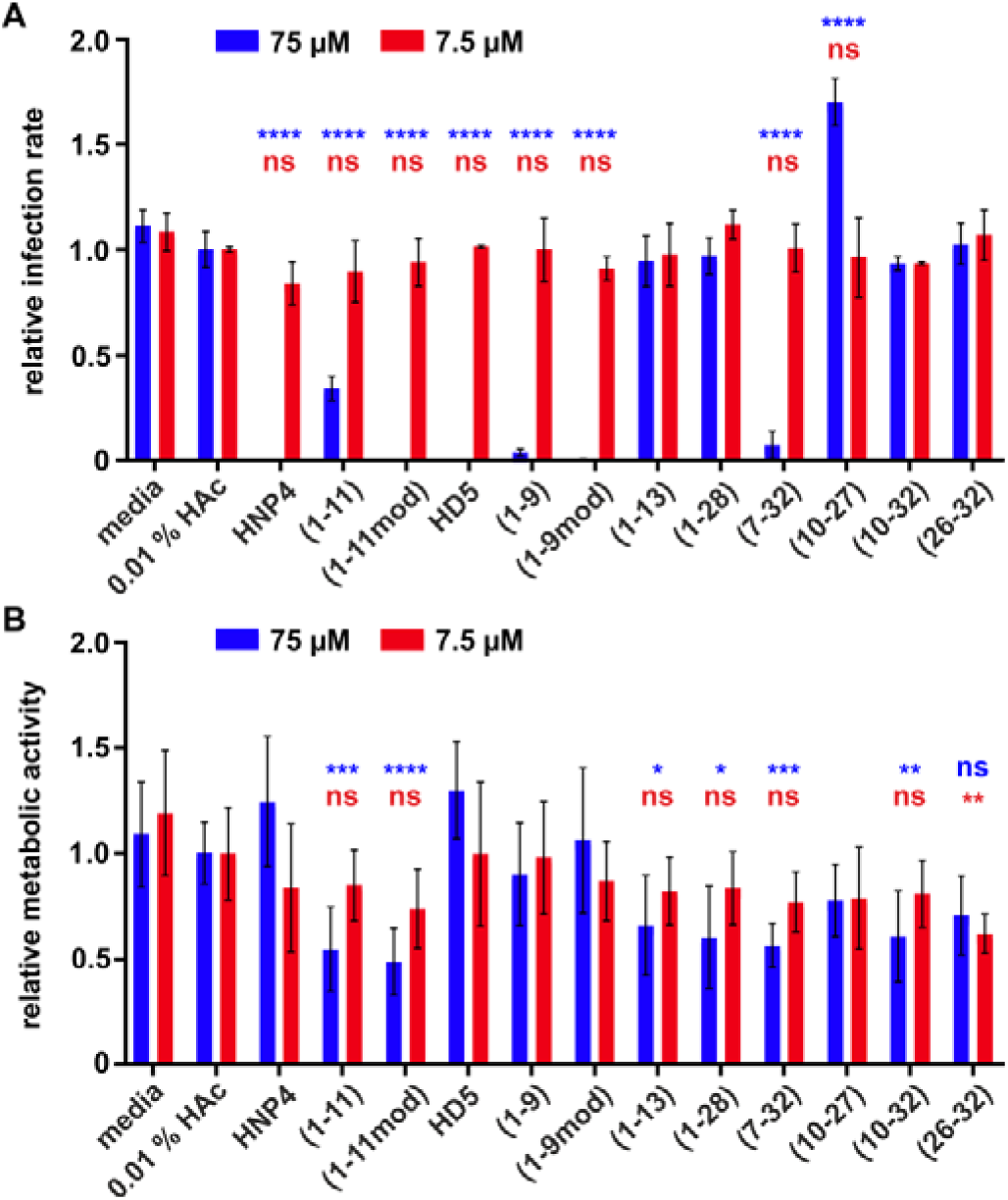
Antiviral activity and cytotoxicity of α-defensin-derived peptides. (A) HFF were infected with TB40/E-ΔUL16-eGFP (MOI of 0.5) and treated with the indicated peptides in concentrations of 7.5 µM and 75 µM. After 40 h of incubation cells were fixed, nuclei stained with DAPI and infection rates measured by imaging with a microplate imager and automated counting of DAPI+ and GFP+ cells. The graph shows the calculated relative infection rate (GFP+/DAPI+, normalized to 0.01 % HAc) **(B)** HFF were treated with the indicated peptides in concentrations of 7.5 µM and 75 µM. After 40 h incubation metabolic activity of cells was measured by MTT. The graph shows the relative metabolic activity (absorption at 570 nm, normalized to 0.01 % HAc). For (A) and (B): both graphs show mean ± SD from triplicate measurements of three independent experiments). ns, not significant; ****, p < 0.0001; ***, p < 0.001; **, p < 0.01; *, p < 0.05. Statistical test used: ordinary one-way-ANOVA with multiple comparisons with Dunnett correction.

To evaluate potential toxicity of the peptides, we incubated HFF cells for 40 hours with the respective peptides and measured viability via MTT (Fig. 2B). This revealed moderate impairment of viability upon incubation with high concentrations of HNP4(1-11), HNP4(1-11mod) and all HD5-fragments with the exception of HD5(1-9) and HD5(1-9mod). Hence, antiviral effects exerted by HNP4, HD5 as well as HD5(1-9) and the modified version are not due to potential toxic effects of these peptides that would impair cellular viability. To further corroborate this finding, we visually inspected HCMV-infected and peptide-treated cells by fluorescence microscopy and identified cells by DAPI-staining (Fig. 3). HCMV-infected cells express GFP as infection marker. Medium or solvent-treated cells show evenly distributed DAPI dots, resembling the nuclei of the cells within the monolayer. This pattern changes upon HCMV-infection, now showing several syncytia-like giant GFP+ cells, which is a cytopathic effect observed upon HCMV-infection (23). Addition of the antiviral active defensins, e.g. HNP4, HD5 or HD5(1-9), but not HD5(1-13) as an example of an inactive peptide, completely reverted this phenotype, blocked HCMV-infection as evident by the reduction of GFP+ cells and restored the homogenous morphology of the cellular monolayer, thus preventing HCMV-induced cytopathic effects (Fig. 3). In conclusion, we identified HNP4, HD5 as well as the HD5-derived peptide HD5(1-9) as natural human peptide inhibitors of HCMV.

**Figure 3:**
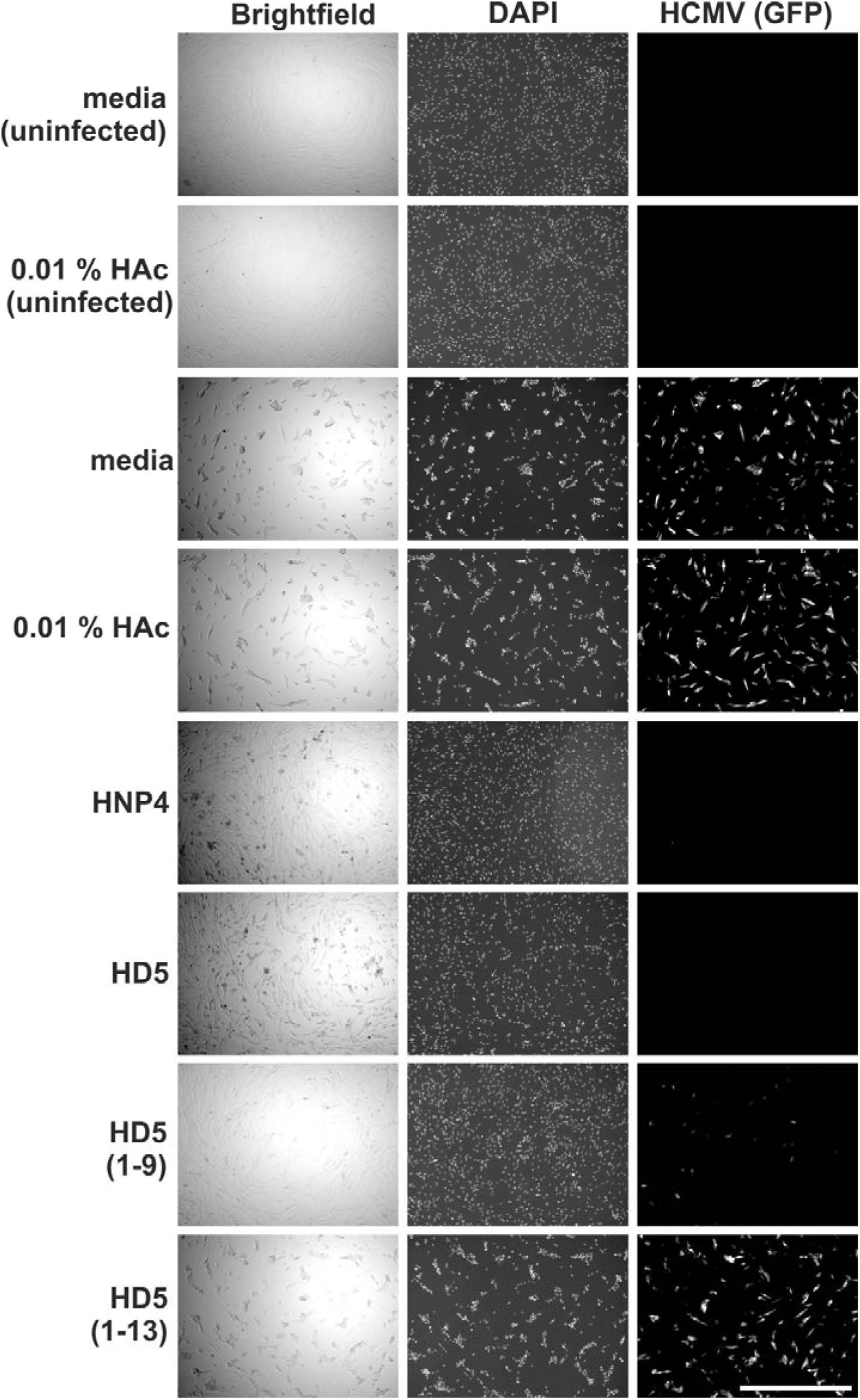
Suppression of HCMV infection and cytopathic effects by α-defensin-derived peptides. Representative images of HFF cells treated with 75 µM of peptide as detailed in Fig. 2A. The scale bar indicates 1000 µm.

### Cytotoxic potential exerted by α-defensin-derived peptides

Our initial assessment of peptide toxicity was after 40 hours of incubation by MTT, hence measuring mitochondrial activity of NADH/NADPH (24). To more carefully evaluate potential toxic effects of our α-defensin peptides, we decided to use xCELLigence real-time monitoring of cellular viability (25). This system continuously measures the electrical impedance of single microtiter-plate wells. Conceivably, upon cell growth, impedance increases, whereas growth arrest or cellular detachment due to dying cells result in a drop of impedance. We further decided to test our collection of peptides not only on HFF, but also on ARPE-19 as a model cell line for epithelial cells and differentiated THP-1 or primary macrophages to model myeloid cells. All of these cells represent important HCMV target cell types *in vivo* (26, 27). Based on their ability to inhibit HCMV-infection at 75 µM (Fig. 2A), we analyzed HNP4, HNP4(1-11), HNP4(1-11mod), HD5, HD5(1-9), HD5(1-9mod) and HD5(7-32) (Fig. 4). Cells were first allowed to adhere. 24 hours post seeding we added the peptides at the concentrations indicated, and impedance was measured every 30 minutes. As expected, all effects observed were dose-dependent. HNP4 started to inhibit growth of all cell types at concentrations of 10 µM and was clearly cytotoxic at higher concentrations. HNP4(1-11) was only toxic when used above 75 µM in HFF and ARPE-19 cells and non-toxic in primary macrophages. HNP4(1-11mod) induced cell death in all cell types starting at around 37.5 µM. Similarly, HD5 showed high cytotoxic potential and already affected growth of for instance primary macrophages at 2-5 µM. In strict contrast, HD5(1-9) was not cytotoxic for any of the cell types tested and only slightly affected growth of HFF and ARPE-19 at 150 µM (Fig. 4). HD5(1-9mod) clearly reduced cell viability from 18.75 µM on, whereas the longer peptide HD5(7-32) only showed cytotoxicity at very high concentrations. In sum, of all defensin peptides tested, only HD5(1-9) did not impair viability of the three different cell types over prolonged incubation.

**Figure 4:**
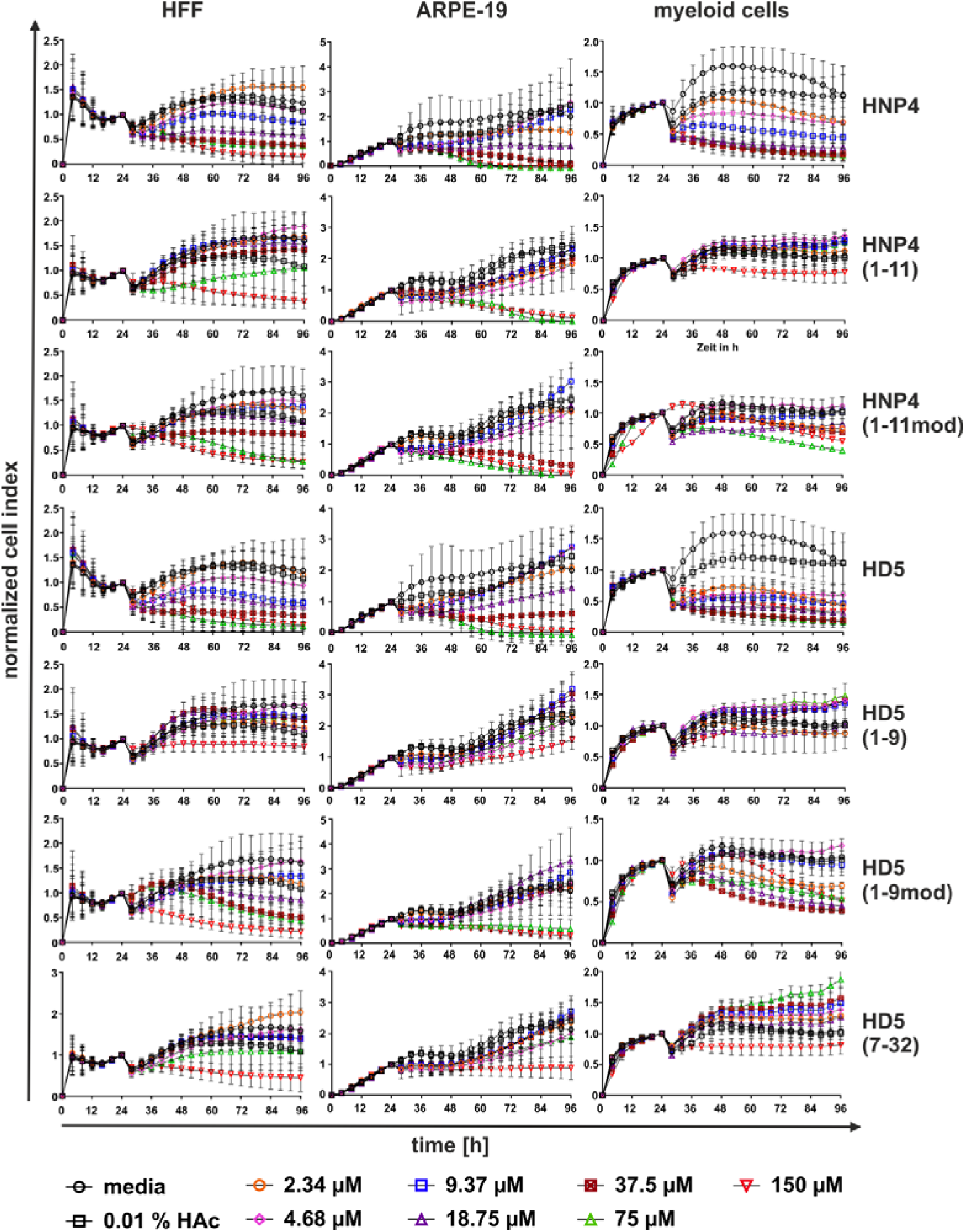
Impact of α-defensin-derived peptides on cellular attachment and growth. HFF, ARPE-19 and macrophages were treated 24 h post seeding with the indicated concentrations of the different peptides. Measurement of electrical impedance was done every 30 min over 72 h (normalized to 24 h value, mean ± SD from duplicates of three independent measurements for HFF and ARPE-19 and one to three independent measurements for macrophages).

### Dose-dependent inhibition of HCMV-infection by α-defensin-derived peptides

Next, we analyzed the anti-HCMV activity of the selected candidate peptides on the three different cell lines in a dose-dependent manner. This allows calculating the inhibitory concentration 50 (IC50), i.e. the drug-concentration at which half-maximal antiviral activity is achieved. Again, cells were infected with HCMV at an MOI of 0.5 and the peptides were added together with the virus inoculum. 40 hours later cells were fixed and stained with DAPI and the viral immediate early (IE1/2) protein. Then infection rate was quantified by automated fluorescence microscopy (Fig. 5). In agreement with our previous data (Fig. 2A), all defensins inhibited HCMV infection in a dose-dependent manner with only slight differences among the cell lines tested. However, since HNP4, HNP4(1-11mod), HD5 and HD5(1-9mod) also affected cell viability in our xCELLigence measurements (Fig. 4), these results have to be interpreted very carefully. Overall, as before, only HD5(1-9) showed antiviral activity at medium (50 µM) to high concentrations (100 µM) without cytotoxic effects (Fig.4 and Fig. 5). This is even more evident by looking at the CC50 (i.e. the drug-concentration at which half-maximal cytotoxic activity is observed) and IC50 values calculated from the data obtained from ARPE-19 cells as an example (Fig. 6). CC50 (Fig. 6A) and IC50 (Fig. 6B) values of HNP4, HNP4(1-11) and HD5 were highly similar, indicating that these peptides do not exert specific antiviral activity. HNP4(1-11mod) and HD5(7-32) are about two- to three-fold less toxic than antiviral, and HD5(1-9mod) was antiviral with an IC50 of ∼14 µM and clearly cytotoxic with a CC50 of ∼57 µM. Finally, HD5(1-9) had an IC50 of ∼40 µM and CC50 > 150 µM indicating that this HD5-derived peptide fragment exerts specific antiviral activity against HCMV.

**Figure 5:**
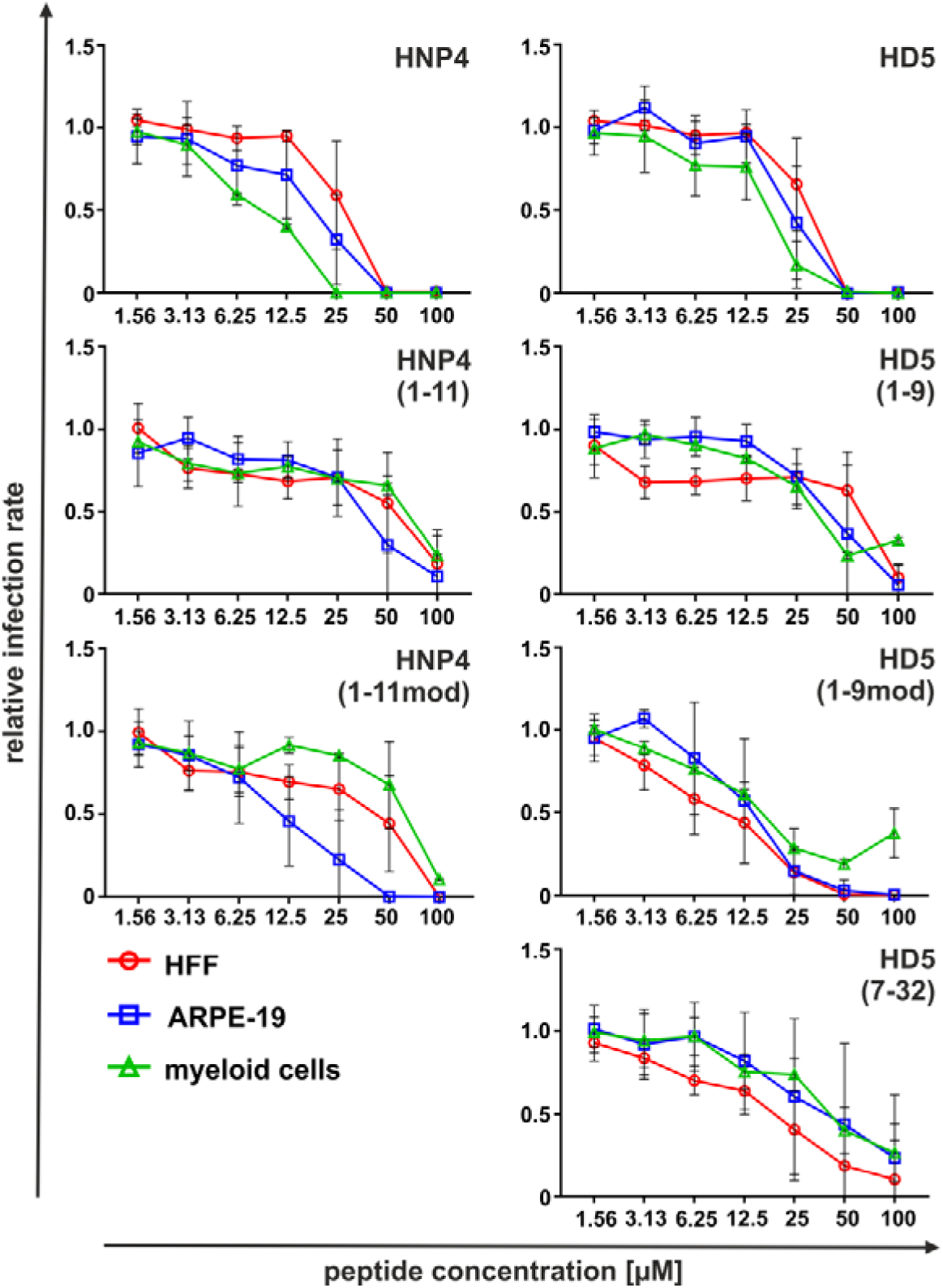
Dose-dependent inhibition of HCMV-infection by α-defensin-derived peptides. HFF, ARPE-19 and PMA-differentiated THP-1 cells or macrophages were infected with HCMV TB40/E-ΔUL16-EGFP and treated with the various peptides in different concentrations. After 40 h incubation, cells were fixed and HCMV-infected cells identified by IE1/2 antigen staining (in addition to GFP staining we used IE1/2 for higher sensitivity as compared to the experiment shown in Fig. 1). Nuclear staining was done with DAPI. Infection rates were measured by imaging with a microplate imager and automated counting of DAPI+ and IE1/2+ cells. The calculated infection rate for all three cell types is shown (IE1/2+/DAPI+, normalized to medium only, mean ± SD from duplicate infections of three independent experiments for HFF and ARPE-19 and one experiment for THP-1/macrophages).

**Figure 6:**
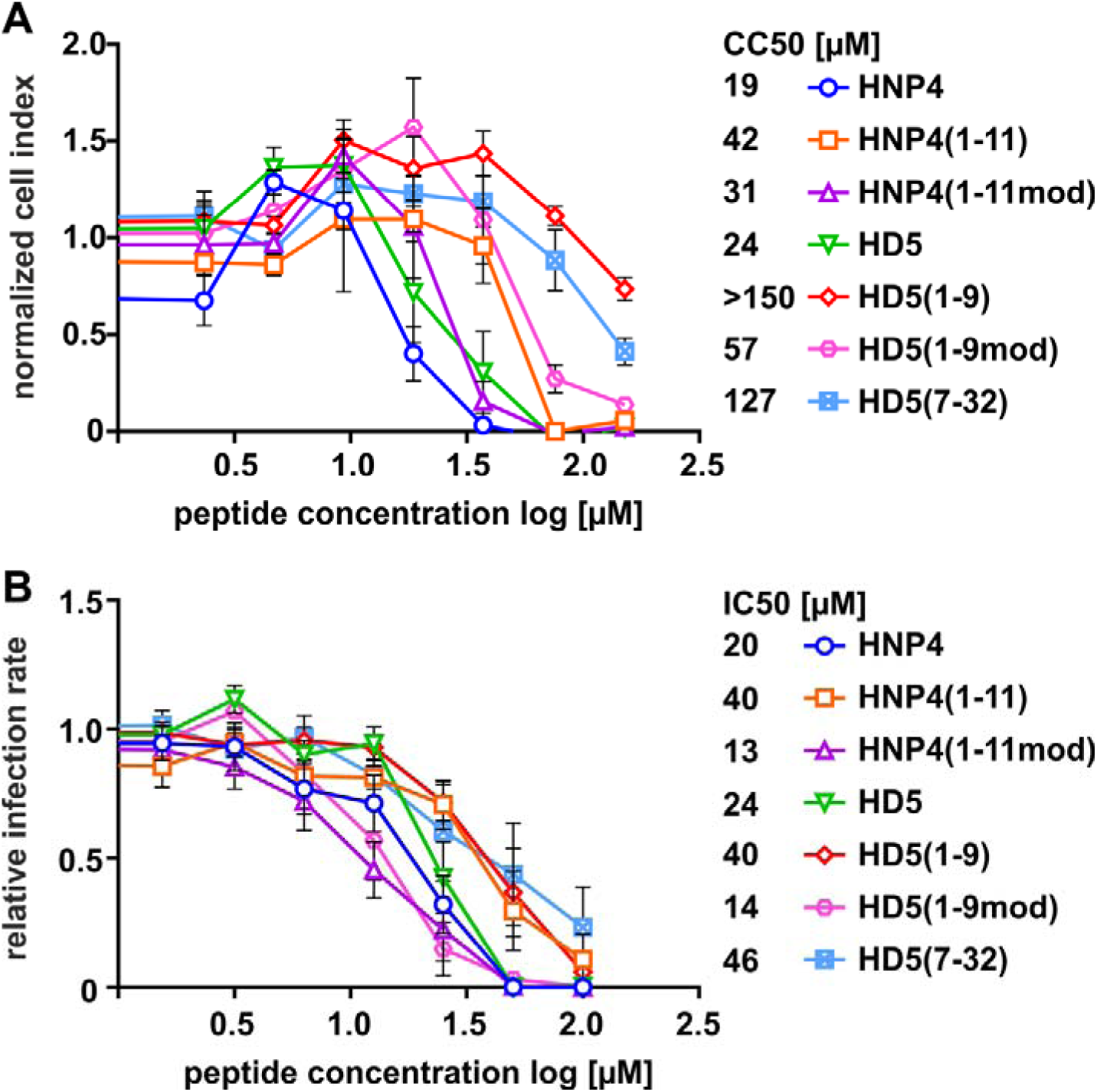
Calculation of IC50 and CC50 for α-defensin-derived peptides. **(A)** CC50 was determined by measuring electrical impedance of ARPE-19 cells at 72 h post incubation (96 h total time) with different concentrations of the respective peptides (compare Fig. 4). **(B)** IC50 was determined by measuring HCMV infection of ARPE-19 cells post incubation with different concentrations of the respective peptides (compare Fig. 4). Mean ± SEM from three independent experiments with duplicate infections.

### Effect of HD5(1-9) on embryonic development

As a first test of HD5(1-9) toxicity *in vivo,* as well as to elucidate potential effects of the peptide on embryonic development, experiments with zebrafish embryos were performed (28). Embryos were treated with HD5(1-9) at concentrations of 25 µM, 75 µM, 125 µM and 250 µM starting at 6-7 hours post fertilization (hpf), which is the time point recommended by pharmaceutical company-utilized assays to assess toxic effects of compounds on early embryonic development (29). Phenotypes were analyzed at 13, 24 and 48 hpf (Fig. 7). At the highest peptide concentration of 250 µM, half of the embryos had died by 13 hpf, indicating that excess of HD5(1-9) can affect embryonic development. Impaired development at 250 µM peptide exposure was also evident over the whole observation period, leading to the death of 11 embryos and 4 embryos having severe developmental delays by 48 hpf (compare Fig. 7B, 7C and examples in Fig. 7D). Reduction of the peptide concentration to 125 µM reduced the negative effects on embryonic development, with approximately half of the embryos having no alterations (Fig. 7C and 7D). Furthermore, all embryos treated with 25 µM or 75 µM of HD5(1-9) developed normally, except for one embryo that had died by 13 hpf due to unknown reasons (Fig. 7). Altogether, HD5(1-9) concentrations of 25 µM to 75 µM, which are higher than the IC50 of ∼40 µM, do not affect zebrafish embryonic development or show toxicity *in vivo*.

**Figure 7:**
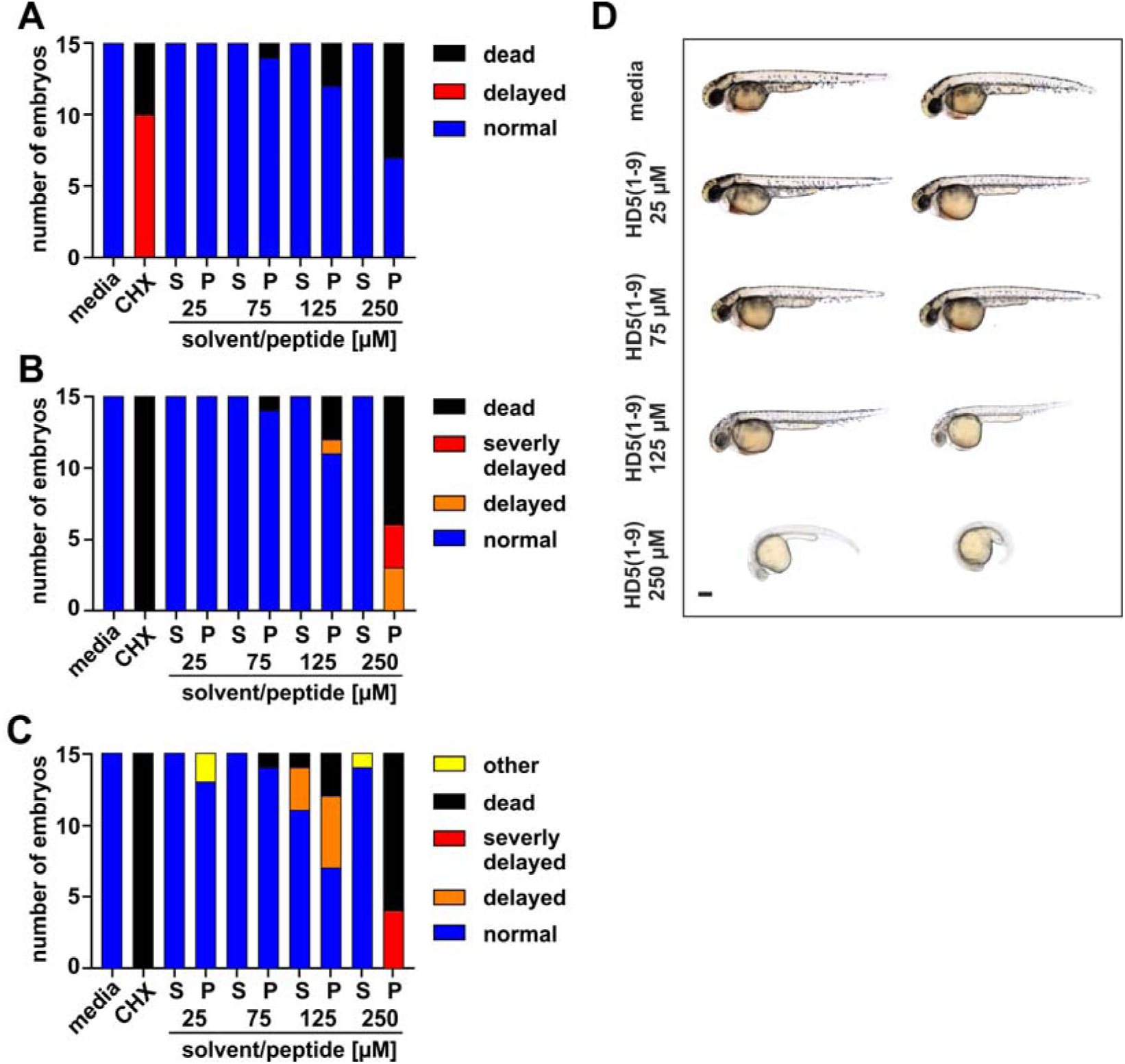
Impact of HD5(1-9) on embryonic development of zebrafish. 6-7 h post-fertilization (hpf), peptides at different concentrations were added to the embryos. At 13 hpf (A), 24 hpf (B) and 48 hpf (C) the embryos were visually inspected. Solvent: PBS, CHX: cycloheximide. (D) Representative images of phenotypes at 48 hpf are shown. The scale bar indicates 250 µM.

### HD5(1-9) inhibits multiresistant, primary HCMV isolates

HCMV TB40/E is a lab-adapted strain that might differ from primary HCMV isolates in terms of cellular tropism, infectivity or cytopathic properties. We hence tested whether HD5(1-9) is also active against primary HCMV isolates from different compartments of patients including amniotic fluid, breast milk and leukocytes (Fig. 8). Of note, the leukocyte isolate from a stem cell transplant recipient is genotypically (mutations UL97 L595S, UL54 V715M) and phenotypically resistant against GCV (IC50 > 30 µM), PFA (IC50 = 795 µM), and CDV (IC50 = 1.8 µM) (4), while both isolates from amniotic fluid and breast milk were therapy-naive. We infected HFF cells at MOIs of 0.2 and 0.1 in the presence of increasing amounts of HD5(1-9). Similar to our previous results, HD5(1-9) inhibited HCMV infection efficiently at a concentration of 100 μM in HFF. Importantly, not only infection with the lab-adapted TB40/E strain was blocked, but also infection with the primary isolates was sensitive towards inhibition by HD5(1-9). Hence, HD5(1-9) is a peptide inhibitor active against primary as well as multiresistant HCMV strains.

**Figure 8:**
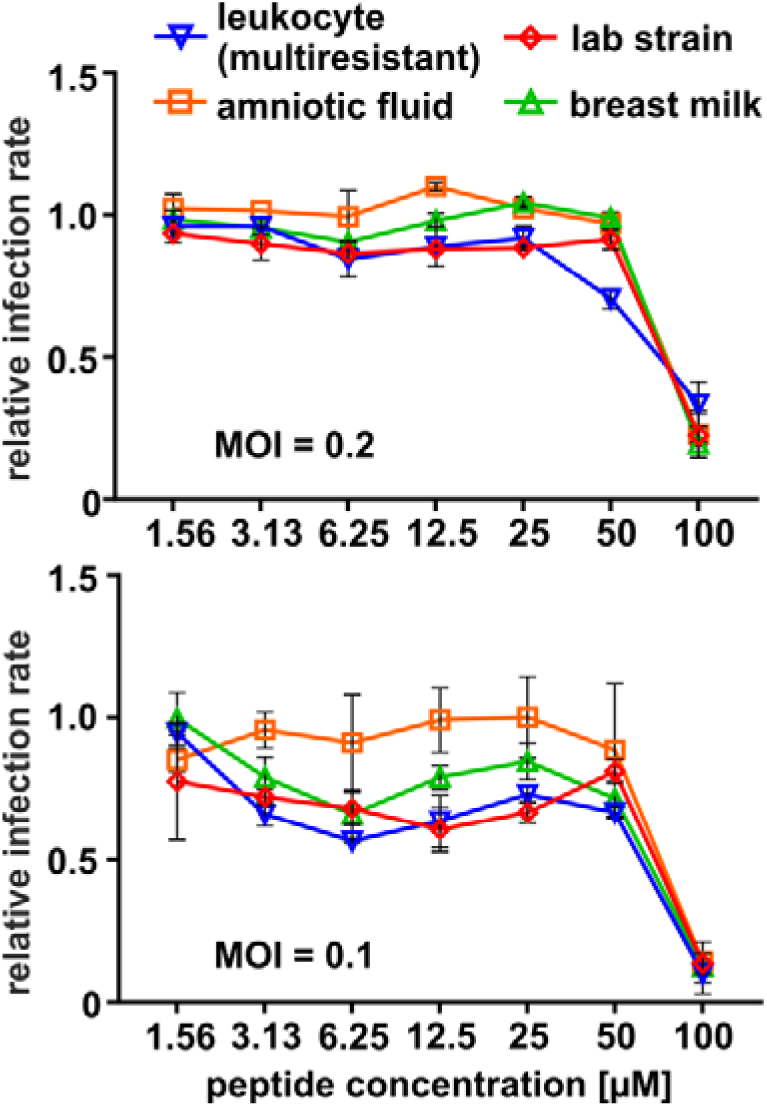
HD5(1-9) inhibits infection with primary patient-derived HCMV isolates. HFF were infected with different clinical HCMV isolates and TB40/E-ΔUL16-eGFP as reference strain with an MOI of 0.2 or 0.1 and treated with HD5(1-9) in different concentrations. After 40 h incubation, cells were fixed and HCMV-infected cells identified by IE1/2 antigen staining. Nuclear staining was done with DAPI. Infection rates were measured by imaging with a microplate imager and automated counting of DAPI+ and IE1/2+ cells. The graphs show the calculated infection rate (IE1/2+/DAPI+, normalized to medium only, mean ± SD from duplicate infections each).

### Structure-activity relationship of HD5(1-9) and HD5(1-13)

In order to identify relevant amino acids regarding antiviral activity, HD5-derived peptides were designed. Arginine (Arg) residues were suggested to participate in HD5-membrane interactions, and hence HD5-mediated antimicrobial effects (30). Furthermore, Cysteine (Cys) at positions three and five are involved in forming disulfide bridges in the context of the full length protein and contribute to the biological activity of defensins (7, 16, 31). Therefore, we ordered three HD5(1-9) modified peptides: HD5(1-9)[R>A], where two Arg residues were mutated to alanine (Ala); HD5(1-9)[C>S] where both Cys were mutated to serine (Ser) and HD5(1-9)[C3S; C5R] where Cys3 was mutated to Ser and Cys5 to Arg. The latter one was done to test the effect of a membrane-interaction promoting additional Arg instead of a Cys (compare with HD5(1-9) peptide structure depicted in Fig.9A).

**Figure 9:**
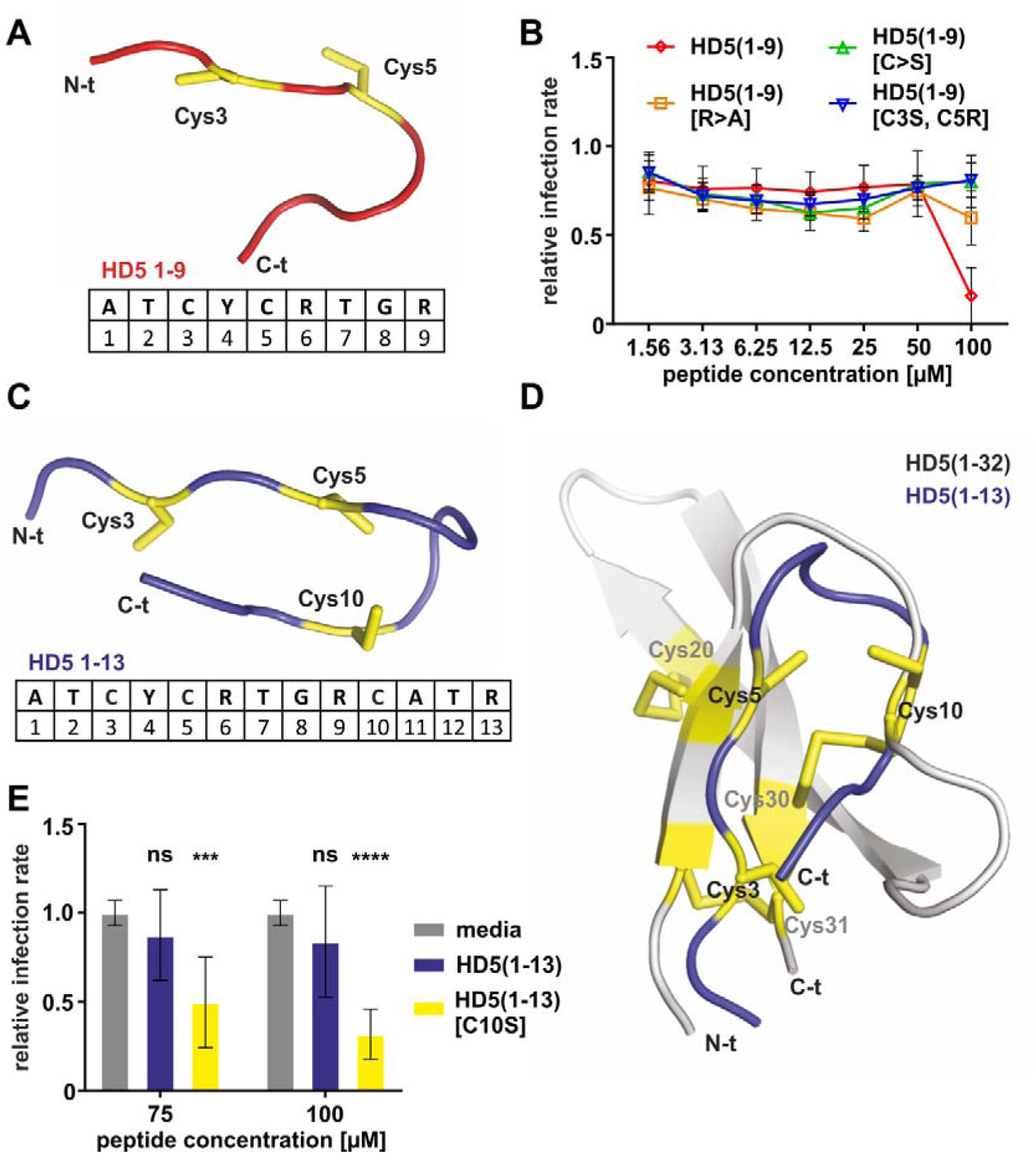
Cysteine and arginine residues in HD5(1-9) mediate its antiviral activity. **(A)** Model and aa-sequence of HD5(1-9); structural model generated by PEP-FOLD 3 (32). **(B)** HFF were infected with TB40/E-ΔUL16-eGFP (MOI of 0.2) and treated with the indicated derivatives of the peptide fragment HD5(1-9) in different concentrations. After 40 h incubation, cells were fixed and HCMV-infected cells identified by IE1/2 antigen staining. Nuclear staining was done with DAPI. Infection rates were measured by imaging with a microplate imager and automated counting of DAPI+ and IE1/2+ cells. The graphs show the calculated infection rate (IE1/2+/DAPI+, normalized to medium only, mean ± SD from duplicate infections of three independent experiments). **(C)** Model and aa-sequence of HD5(1-13); structural model generated by PEP-FOLD 3. **(D)** Structural alignment of HD5(1-32) crystal structure (39) (pdb: 2lxz) (grey), and prediction model of peptide HD5(1-13) generated by PEP FOLD 3 (blue). Cys are marked in yellow and arrows represent β-strands. Alignment generated with PyMol. **(E)** Similar experimental setup as in (B), however HD5(1-13) and HD5(1-13)[C10S] with a mutated cysteine at position ten to serine were used (mean ± SD from duplicate infections of three independent experiments). ns, not significant; ****, p < 0.0001; ***, p < 0.001. Statistical test used: ordinary one-way-ANOVA with multiple comparisons with Dunnett correction.

As before, HD5(1-9) inhibited HCMV at concentrations above 50 µM when infecting HFF. In contrast, all of the modifications introduced abrogated the antiviral activity of HD5(1-9) (Fig. 9B). In conclusion, Cys as well as Arg residues within HD5(1-9) are important for the full antiviral activity of the peptide.

In our initial screen HD5(1-9) showed antiviral activity, whereas the four amino acid longer version HD5(1-13) was inactive (Fig. 2A). The peptide HD5(1-9) was predicted to be flexible and extended in solution (Fig. 9A), which was also confirmed by NMR spectroscopy. In contrast, algorithms to predict peptide structures (PEP-FOLD3, (32)) show that HD5(1-13) is likely to adopt a “close” conformation, which might be stabilized by the formation of a disulfide bond between Cys5 and Cys10 (Fig. 9C and D). This close conformation could explain the difference in activity between the peptides. To challenge this hypothesis, we designed and ordered another peptide HD5(1-13)[C10S], with the goal of opening the peptide conformation by disrupting a potential disulfide bridge formation involving Cys10. In keeping with previous results, HD5(1-13) did not inhibit HCMV infection (Fig. 9E). In contrast, HD5(1-13)[C10S] showed a gain-of-function phenotype and blocked HCMV infection at 75 µM and 100 µM.

Together, the data demonstrates that the antiviral activity of HD5-derived peptides is specific and can be enhanced by distinct amino acid modifications.

### HD5(1-9) interferes with HCMV attachment and entry

In our infection assays, GFP controlled by the UL16 promoter or immunostaining of the HCMV IE1/2 antigen are markers for early viral gene expression, before *de novo* viral genome replication. We hence hypothesize that HD5(1-9) blocks an early event in the HCMV life cycle. We first set up an experiment to elucidate if HD5(1-9) acts directly on viral particles or exerts its antiviral activity on a cellular basis. For this, we pre-incubated virus stocks used for infection in small volumes for 1 h at 37 °C with the indicated concentrations of the peptide and added the mixture then to HFF, or performed the experiment as before, i.e. adding virus and peptide simultaneously to the cells (Fig. 10A). In one condition, to assess if concentrated peptide is sufficient to neutralize infectivity of viral particles, HCMV stock and peptide were pre-incubated at 100 µM of HD5(1-9). The mixture was then diluted 10-fold upon addition to the cells, resulting in a final concentration of 10 µM HD5(1-9) during infection. However, neither pre-incubation of the peptide with virus stocks, nor incubation of virus in concentrated peptide solution increased antiviral activity (Fig. 10A). These results suggest that HD5(1-9) exerts its antiviral activity not on assembled viral particles, but in the context of the cellular infection process.

**Figure 10:**
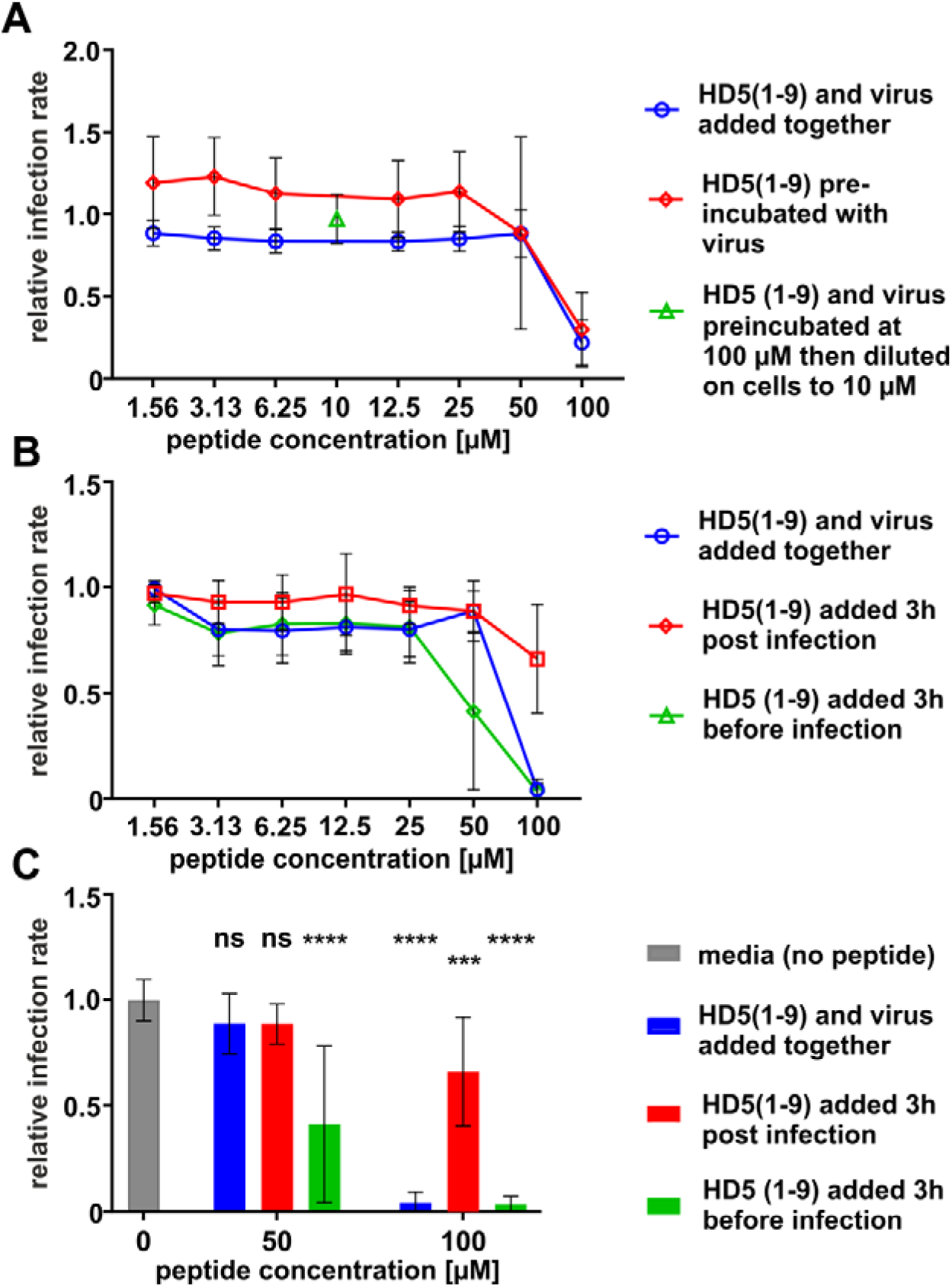
HD5(1-9) inhibits an early step during HCMV entry. **(A)** HFF were infected with TB40/E-ΔUL16-eGFP at a MOI of 0.2. HD5(1-9) was either given directly to the cells together with the viral inoculum or pre-incubated for 1 h at 37 °C with the virus preparation and then added to the cells. In one condition, HD5(1-9) and the virus preparation were pre-incubated at a concentration of 100 µM and diluted ten-fold on the cells to a final HD5(1-9) concentration of 10 µM. After 40 h incubation, cells were fixed and HCMV-infected cells identified by IE1/2 antigen staining. Nuclear staining was done with DAPI. Infection rates were measured by imaging with a microplate imager and automated counting of DAPI+ and IE1/2+ cells. The graph shows the calculated infection rate (IE1/2+/DAPI+, normalized to medium only, mean ± SD from duplicate infections of three independent experiments). **(B)** Similar infection protocol as in (A) however we tested two conditions in which HD5(1-9) was given to the cells either 3 hours before, or alternatively 3 hours post infection with TB40/E-ΔUL16-eGFP. The graph shows the calculated infection rate (IE1/2+/DAPI+, normalized to medium only, mean ± SD from duplicate infections of three independent experiments). **(C)** Statistical analysis of the data depicted in (B) for concentrations of 50 µM and 100 µM peptide. ns, not significant; ****, p < 0.0001; ***, p < 0.001. Statistical test used: ordinary one-way-ANOVA with multiple comparisons with Dunnett correction.

We next used time-of-addition assays to investigate whether HD5(1-9) inhibits a pre- or post-entry step of HCMV infection. For this, we added HD5(1-9) either (i) three hours before infection, (ii) with the virus inoculum or (iii) three hours post-infection to the cells (Fig 10B). Of note, when HD5(1-9) was added to the cells three hours before infection, its activity was markedly increased. To the contrary, when added three hours post-infection, the peptide was nearly inactive in blocking HCMV-infection, even at a high concentration of 100 µM (Fig 10C). This indicates that the antiviral activity of HD5(1-9) is attributable to an inhibition of HCMV cellular attachment or entry.

To directly asses, if HD5(1-9) blocks attachment of HCMV particles or entry of cell surface bound viruses, we employed a dual-fluorescently labeled virus (33). This allows to discriminate enveloped surface bound viruses (GFP+/mCherry+), from particles having lost their mCherry-labeled envelope during entry now appearing GFP+ only. We infected HFF cells at an MOI of 2 and added HD5(1-9) either during infection or 3 h later. Then cells were fixed, stained with Hoechst33342 and the amount of cell surface bound (GFP+/mCherry+) as well as penetrated viral particles (GFP+) quantified by fluorescence microscopy (Fig. 11). Upon addition of HD5(1-9), but not the inactive mutated peptide HD5(1-9)[R>A], the number of total cell surface bound particles was strongly reduced (quantitative analysis in Fig. 11A and representative images compare Fig. 11B). In contrast, when the same assay was performed with peptide added 3 h post infection, HD5(1-9) did not have an inhibitory effect on the amount of cell surface bound or cell penetrated particles, which is overall consistent with an inhibition of HCMV attachment by HD5(1-9).

**Figure 11:**
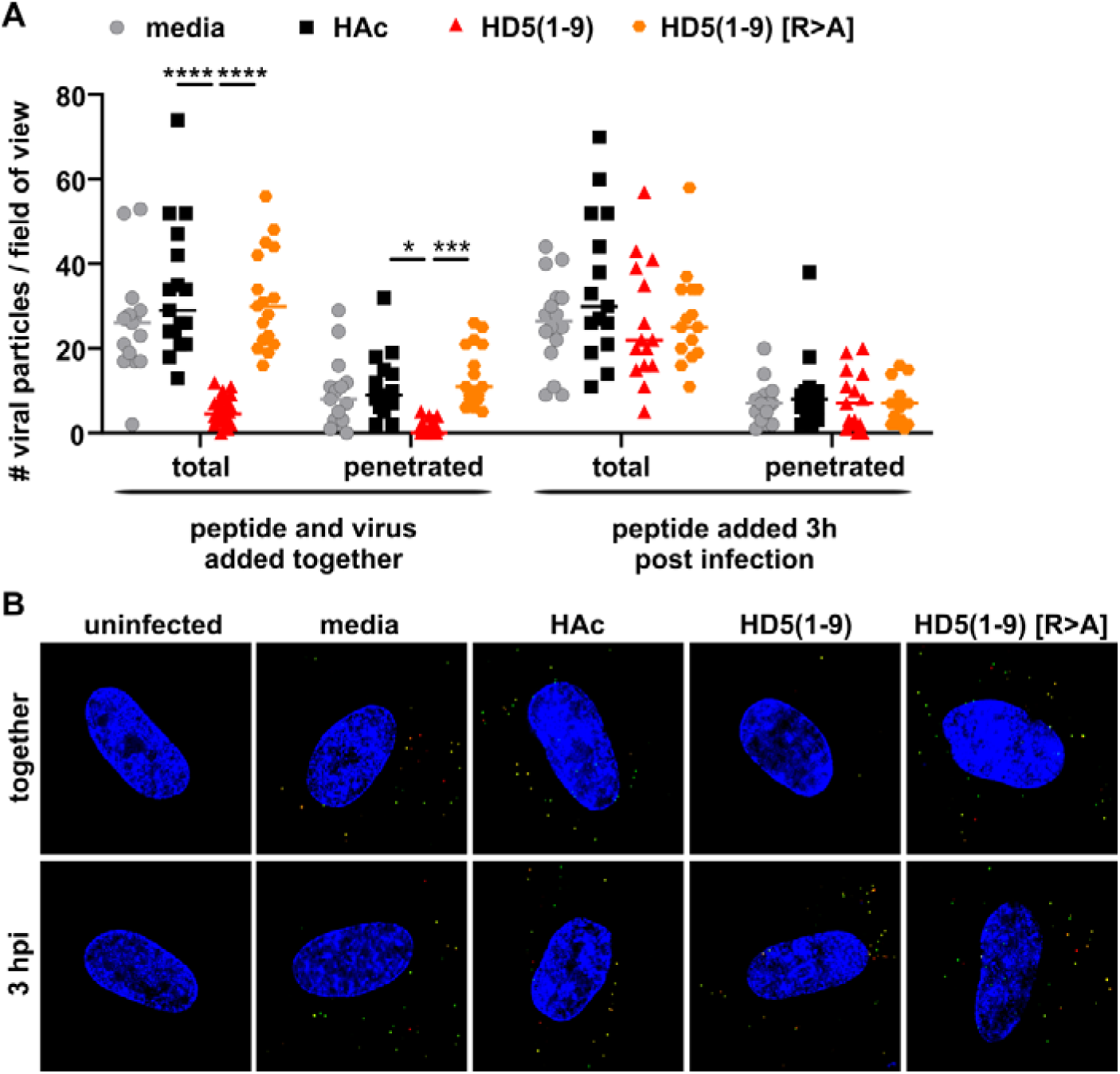
HD5(1-9) inhibits cellular attachment of HCMV particles. HFF were infected at an MOI of 2 with the dual fluorescent virus TB40-BAC_KL_-UL32eGFP-UL100mCherry, expressing pp150 (pUL32) fused to eGFP and gM (pUL100) fused to mCherry. This allows to discriminate enveloped virus particles (eGFP+/mCherry+), from penetrated viruses (eGFP+). HD5(1-9) or HD5(1-9) [R>A] were added either simultaneously with the virus or 3 hpi at a final concentration of 100 µM. 6 h later cells were fixed and DNA stained and subjected to fluorescence microscopy. (A) For each condition, the total number of cell-associated viral particles or penetrated particles was counted for at least 15 cells. Statistical test used: ordinary one-way-ANOVA with multiple comparisons with Dunnett correction; ****, p < 0.0001; ***, p < 0.001; *, p < 0.05. (B) Representative images of the different conditions quantified in (A). The result was confirmed in an additional independent experiment.

## DISCUSSION

We here identified a natural human α-defensin HD5-derived peptide as attachment inhibitor of HCMV. HD5(1-9) was non-toxic up to concentrations of 150 µM for diverse cell types and showed antiviral activity starting at concentrations higher than 50 µM in HFF cells and 25 µM in ARPE-19 with an IC50 of ∼40 µM in ARPE-19 (Fig. 6). This antiviral effect was highly specific, since we could disrupt the activity of the peptide by mutating Cys3 and Cys5 as well as Arg6 and Arg9 (Fig. 9). In addition, other HD5-derived peptides did not show antiviral activity or were clearly cytotoxic when used at similar concentrations. Of note, we could also demonstrate a gain-of-function for the four amino acid longer version HD5(1-13) by mutation of Cys10 to Ser. This could be explained by the opening of the close conformation predicted for HD5(1-13), since the substitution of Cys10 to Ser would impair the formation of a disulfide bridge, which could maintain the structure closed. This confers antiviral activity to HD5(1-13) blocking HCMV infection at 75 µM and above (Fig. 9). Moreover, this demonstrates that it is possible to improve antiviral activity of the small HD5-derived peptides *per se* by single amino acid exchanges. Hence, it is tempting to speculate that the activity of HD5(1-9) could be improved for instance by shortening of the N-terminus or changing glycine at position eight into arginine to further increase the positive charge of the peptide. However, such modifications have to be introduced with caution. HD5(1-9mod), composed of D-amino acids and equipped with an N-terminal acetate and a C-terminal amide-group to render it more stable and prevent degradation, showed high toxicity over prolonged incubation on various cell types (Fig. 4).

In this context, it has to be noted that toxicity is a critical parameter when determining the antiviral activity of drug candidates. Viruses are obligatory intracellular pathogens that use the cellular machinery for propagation. Hence, any compound-induced impairment of viability could negatively impact viral replication. We performed various independent assays to determine the potential cytotoxicity of our peptide candidates: (i) MTT assays measuring the metabolic activity of cells; (ii) adherence and growth by monitoring the electrical impedance of cells when they are cultured as monolayers on plates; and (iii) peptide effects on embryonic development of zebrafish. Remarkably, while in MTT none of the peptides dramatically affected metabolic activity (Fig. 2B), only HD5(1-9) did not impair long-term growth and adherence of cells in high concentration (Fig. 4). In zebrafish embryonic development, we did not observe toxicity of HD5(1-9) at concentrations that were nearly twice as high as its IC50 (Fig. 7).

Defensins have been described as antiviral effector molecules of various viruses with different potential modes of action (16, 34). Nevertheless, this is the first comprehensive report demonstrating antiviral activity of a defensin against HCMV and in particular of the α-defensin HD5-derived peptide HD5(1-9). Just recently, Ehmann and colleagues provided compelling evidence that defensins are cleaved by proteolytic processes and that the resulting peptides - including HD5(1-9) - have broad antimicrobial activity (22). Administration of HD5(1-9) via the oral route was well tolerated in mice and elicited microbiome-modulating activity *in vivo*. However, neither parenteral administration nor bioavailability of HD5(1-9) in blood and organs after oral consumption have been analyzed yet.

HCMV poses a serious threat for immunocompromised patients, for instance HIV-1-infected individuals or transplant recipients. Treatment with GCV is often problematic, since it has a high nephrotoxic potential and the resistance barrier is low. The terminase inhibitor Letermovir is a therapeutic alternative, but first resistance-conferring mutations have been described (3, 5). In this regard, the establishment and testing of new treatment options is necessary. Even though the HD5-derived peptides analyzed within our study are at a proof-of-principle stage and currently far from being used as antiviral drugs, their further development holds a variety of potentially attractive promises: (i) due to the small size of just nine amino acids HD5(1-9) is affordable and realistic to develop, even as peptide inhibitor; (ii) the mode of action is block of viral attachment to cells, thereby protecting them from infection; (iii) HD5(1-9) acts on a cellular target, minimizing the risk of resistance development; (iv) as attachment inhibitor, HD5(1-9) binds to the cell surface and hence does not have to penetrate cells; (v) HD5(1-9) has a broad antimicrobial activity and therefore might also protect from bacterial as well as other viral infections.

Altogether, we provide proof-of-concept for the use of α-defensin HD5-derived peptides, in particular HD5(1-9), as potential entry inhibitors of lab-adapted, as well as primary and multiresistant human cytomegalovirus strains. The advantages associated with the use of defensin-derived peptides for the therapy of human viral diseases warrants their further in-depth analyses and preclinical development.

## MATERIAL AND METHODS

### Cell culture

Primary human macrophages (isolated from buffy coat of healthy blood donors, see details below), primary human foreskin fibroblasts (HFF; from ATCC #SCRC-1041), ARPE-19 (from ATCC #CRL-2302), and THP-1 (from the NIH AIDS reagent program #9942) were cultured at 37 °C with 5% CO_2_. Primary human macrophages were prepared and differentiated as follows and maintained in macrophage-medium (RPMI supplemented with 4% human AB serum, 2 mM L-Glutamine, 100 µg/ml penicillin/streptomycin, 1 mM sodium pyruvate, 1x non-essential amino acids and 0.4x MEM vitamins). HFF and ARPE-19 were cultured in DMEM containing 5% FCS as well as 2 mM L-Glutamine and 100 µg/ml penicillin/streptomycin. THP-1 cells were maintained in RPMI containing 0.25 µg/ml puromycin supplemented with 10% FCS, 2 mM L-Glutamine and 100 µg/ml penicillin/streptomycin. For the respective experiments, THP-1 cells were differentiated with 30 ng/ml phorbol-myristate-acetate (PMA) for 24 h at 37 °C.

### Isolation and differentiation of primary human macrophages

Macrophages were generated from buffy coats of healthy blood donors who gave informed consent for the use of blood-derived products for research purposes. We do not collect data concerning age, gender or ethnicity and comply with all relevant ethical regulations (IRB #507/2017BO1). All buffy coat donations were received in anonymous form and chosen randomly. PBMCs were isolated from buffy coats by biocoll density gradient centrifugation and differentiated 3 days by plastic adherence in macrophage-medium. After 3 days, non-adherent cells were removed by washing, and the macrophages were further differentiated 4 days with macrophage-medium.

### Infection assays and HCMV viral stocks

For generating HCMV stocks HFF cells were infected with TB40-ΔUL16-eGFP essentially as described before (35). Infectious supernatant was harvested 5 to 7 dpi and subsequently cleared from cells and cellular debris by centrifuging 10 min at 3200 x g.

HFF and ARPE-19 were seeded with 10 000, macrophages with 20 000 and THP-1 with 50 000 cells per well of a 96-well plate. Peptides and the virus were added simultaneously. The respective MOI is indicated in the figure legends. 40 hpi, the cells were fixed with 2% PFA (10 min at 37 °C, 20 min at room temperature or overnight at 4 °C) and permeabilized with ice-cold 90 % methanol in H_2_O for 20 min at 4 °C. Cells were further stained for IE1/2 (mouse anti HCMV IE E13, Argene, 1:1000 dilution in PBS) and counterstained with goat-anti-mouse Alexa594 (Thermo, 1:2000 dilution in PBS) followed by DAPI. Images were taken with the Biotek Cytation 3 multiplate reader. The infection rate was calculated by the number of IE1/2-positive signals to the number of DAPI-positive cell nuclei.

### Screening of the peptide set for antiviral activity

Peptides were solved in PBS or 0.01% HAc. The full-length peptide HNP4 and its derivatives HNP4(1-11) and HNP4(1-11mod) as well as the full-length peptide HD5 and its derivatives HD5(1-9), HD5(1-9mod), HD5(1-13), HD5(1-28), HD5(7-32), HD5(10-27), HD5(10-32) and HD5(26-32) were obtained from EMC microcollections GmbH and tested for their antiviral activity against HCMV TB40/E-ΔUL16-eGFP on HFF at concentrations of 7.5 µM and 75 µM respectively. A MOI of 0.5 was used. For evaluation of infection, GFP was used as readout and images were taken with the Biotek Cytation 3 multiplate reader. The infection rate was calculated by the number of GFP-positive cells to the number of DAPI-positive cell nuclei.

### Screening of the peptide set for cytotoxicity: MTT assay

The cell viability of HFF after treatment with the peptides HNP4, HNP4 (1-11), HNP4 (1-11mod), HD5, HD5 (1-9), HD5 (1 9mod), HD5 (1-13), HD5 (1-28), HD5 (7-32), HD5 (10-27), HD5 (10-32) and HD5 (26-32) was measured at different concentrations by standard MTT measurement (24). For the screening tests 10 000 HFF were seeded in 96-well plates. The peptides were tested at concentrations of 7.5 µM and 75 µM. 40 h post treatment, the medium was changed to 90 µl phenol red free medium, and MTT solution was added and incubated for 3 h. The medium was removed and the crystals were dissolved in 100 µl 0.04 M hydrochloric acid in isopropanol for 10 min. Absorption was measured in the Biotek Cytation 3 multiplate reader at 570 and 650 nm. For evaluation, the mean absorption value measured in empty wells was subtracted from all measured absorption values. To determine the absolute absorption the absorption values of the reference wavelength 650 nm were subtracted from the values at 570 nm. The relative absorption was determined by referring the corresponding value to the mean absorption of the HFF treated with 0.01% HAc. The experiment was performed three times and technical triplicates were used.

### Determination of CC50 for selected peptides: impedance measurement

Analogous to the determination of IC50, the toxicity of the peptides HNP4, HNP4(1-11), HNP4(1-11mod), HD5, HD5(1-9), HD5(1-9mod) and HD5(7-32) was investigated in more detail by determining the peptide concentration at which 50 % of the cells show a cytotoxic effect (CC50). The xCELLigence system was used for this purpose. The principle is based on a measurement of the electrical impedance caused by adhesive cells on the bottom of a 96-well plate equipped with microelectrodes. Cell proliferation causes an increase in impedance due to partial isolation of the electrodes, while events leading to altered cell morphology or cell detachment lead to decreased impedance (25).

The experiments were performed with HFF, ARPE-19 and macrophages. 10 000 HFF or ARPE-19 or 20 000 macrophages per well were seeded. The plate was placed in the xCELLigence, and the impedance was measured at 37 °C and 5% CO_2_ for 24 h every 30 min. The plate was then removed, and a media change and treatment with the peptides were performed. The peptide concentrations corresponded to a two-fold serial dilution from 150 µM to 2.34 µM. The positive control was 10% Triton-X. Afterwards, the impedance at the bottom of the 96-well plate was measured every 30 min for a further 72 h. The calculation of the CC50 was done analogously to the calculation of the IC50 in GraphPad Prism 7.0.

### Determination of IC50 for selected peptides

IC50 values were measured for the peptides HNP4, HNP4(1-11), HNP4(1-11mod), HD5, HD5(1-9), HD5(1-9mod) and HD5(7-32). A two-fold serial dilution of the peptides was done from 100 µM to 1.56 µM. The experiments were performed on HFF and ARPE-19 cells with an MOI of 0.5, on THP-1 cells with an MOI of 10 and on primary macrophages with an MOI of 70. The IC50 was calculated in GraphPad Prism 7.0. For this purpose, a dose-response curve was created with the values of the relative infection rate. The relative infection rate of untreated infected cells was the starting point of the curve, and for technical reason the value “0” was substituted in the logarithmic scale for 10^-10^.

### Experiments with zebrafish

To test HD5(1-9) for toxicity and to assess its effect on embryonic development in zebrafish, embryos obtained from natural crosses of wildtype TE fish were used. After mating, embryos were incubated at 28°C until 6 hours post-fertilization (hpf). Triplicates of five embryos each were then placed in wells of a 96-well plate containing 200 µl of different peptide concentrations. HD5(1-9) dissolved in PBS was diluted in normal embryo medium (250 mg/L Instant Ocean salt, 1 mg/L methylene blue in reverse osmosis water adjusted to pH 7 with NaHCO3) (36). The concentrations tested were 25 µM, 75 µM, 125 µM and 250 µM, and the concentration of PBS solvent was adjusted for each control group. At peptide concentrations of 75 µM and above we observed granulae in some wells, which could represent peptide accumulations. As a control, embryos were additionally incubated in 25 µg/ml cycloheximide (C4859, Sigma-Aldrich). After 13 hpf, 24 hpf and 48 hpf a microscopic phenotype analysis was performed. For each time point embryos were automatically imaged using an ACQUIFER Imaging Machine. For the phenotype analysis at 48 hpf, the larvae were manually dechorionated and anaesthetized with 2% tricaine methanesulfonate (A5040-25G, Sigma-Aldrich). Images were acquired on an Axio Zoom.V16 microscope (ZEISS).

### Infection assays with clinical isolates

The antiviral activity of HD5(1-9) in concentrations from 1.56 µM to 100 µM against infection with clinical HCMV isolates was investigated on HFF. In addition to the laboratory-adapted strain TB40/E-ΔUL16-eGFP, which served as reference strain, a breast milk-derived strain, an amniotic fluid-isolated strain and a multidrug-resistant viral isolate from leukocytes of a recipient after the third stem cell transplantation were used. The therapy-naïve strain from cell free milk whey (H1241-2016) was derived from a mother of a preterm infant 10 weeks postpartum during end of viral reactivation. The amnion fluid derived virus strain (H2497-2011) was isolated following termination of pregnancy based on severe fetal brain damage (Preisetanz, S, Diploma thesis, University of Tuebingen, 2012). The multidrug resistant CMV isolate (H815-2006) is already described (4). This viral isolate showed the canonical UL97 mutation L595S and an UL54 mutation V715M, leading to drug resistance against GCV, IC50 = 31.5 µM, and CDV, IC50 = 795 µM. IC50 value against PFA was 1.8 µM. All viral isolates were primarily HFF-adapted and propagated in vitro with at least 10 passages to get TCID values of cell free viral supernatants of 10^5^ to 10^6^/ml.

### Infection assays with derivatives of peptide fragments

For structure-activity relationship HD5(1-9) derivatives with modified amino acids were ordered from JPT Peptide Technologies GmbH. Peptides were solved in PBS or 0.01% HAc at a concentration of 1 mM. The antiviral activity of HD5(1-9), HD5(1-9) [R6>A], HD5(1-9) [C>S], HD5(1-9) [C3S, C5R] and HD5(1-13) [C10S] was tested in concentrations from 1.56 µM to 100 µM on HFF cells. All experiments were performed three times with an MOI of 0.2 using technical duplicates.

### NMR spectroscopy

The unlabeled peptide HD5(1-9) (JPT Technologies) was dissolved in acetic acid 0.01% to a final concentration of 1mM. All NMR spectra required for chemical shift assignment were acquired on Bruker AVIII-600 spectrometer. All spectra were recorded at 298 K. The NMR data were processed using TopSpin 2.1 (Bruker GmbH), and analyzed with Sparky 3.115 (37). In-Phase COSY was acquired with 4096 and 128 complex points in *t*2 and *t*1, respectively, performing 16 scans per increment (38). The TOCSY experiment was recorded with 4096 (*t*2) x 256 (*t*1) complex points using 16 scans per increment and a relaxation delay of 1.5 s. The NOESY spectrum was acquired on Bruker AVIII-800 using 1024 (*t*2) x 172 (*t*1) complex points using 96 scans per increment and a relaxation delay of 1.5 s. The NOESY spectrum was recorded with a NOE mixing time of 80 ms and the TOCSY spectrum was recorded with a spin lock mixing time of 70 ms. The HMQC-COSY and HMBC spectrum were recorded at a resolution of 1024 (*t*2) × 172 (*t*1) complex points, with 256 scans per increment. The ^13^C-HSQC was recorded at a resolution of 1024 (*t*2) × 128 (*t*1) complex points, using 128 scans per increment.

### Preincubation assays

To assess direct binding of HD5(1-9) to viral particles, the peptide was incubated for one hour at 37 °C with the virus in concentrations of 1.56 µM to 100 µM and then given to HFF. As reference, the infection assay was performed at the same concentrations without preincubation. In one condition, peptide and virus were pre-incubated at a peptide concentration of 100 µM in a volume of 10 µl and diluted on the cells to a concentration of 10 µM in 100 µl volume. The experiment was performed three times with a MOI of 0.2 using technical duplicates.

### Time-of-addition assays

(i) Peptide was pre-incubated with cells for 3 h at 37 °C before addition of the virus. (ii) Peptide and virus were added to HFF simultaneously or (iii) the peptide was added to the cells 3 h after the virus. In all conditions, incubation volumes and peptide concentrations were adjusted to achieve a range of peptide concentrations of 1.56 µM to 100 µM. This experiment was performed three times with a MOI of 0.2 using technical duplicates.

### Infection assays with a dual fluorescent virus

In order to investigate the mode of action of HD5(1-9), an infection assay with the dual fluorescent virus TB40-BAC_KL_-UL32eGFP-UL100mCherry on HFF was performed (33). This is an endotheliotropic HCMV strain expressing pp150 (pUL32) fused to eGFP and gM (pUL100) fused to mCherry. This allows discriminating enveloped virus particles, which have the green capsid and the red envelope and thus appear yellow, from already penetrated virus particles which lost their red envelope and only show eGFP signal. 20 000 HFF were seeded in 8-well-chamber-slides (IBIDI). The cells were infected with an MOI of 2 and the peptides (HD5(1-9) and HD5(1-9)[R>A]) were added either simultaneously with the virus or 3 hpi at a final concentration of 100 µM. After 6 h the cells were fixed and stained with 1 µg/ml Hoechst33342 in PBS for 10 min at RT. After fixation and staining, the cells were washed three times with 200 µl PBS each. Images were taken with the Deltavision OMX (GE Healthcare) in confocal mode and virus particles were counted with Fiji (ImageJ).

### Software, analysis tools, and molecular modeling

GraphPad Prism 7.0 was used for statistical analyses and generation of diagrams. The respective statistical test used is indicated in each figure legend. Other software used was Gen5 V2.09 (Biotek), Imaris and SoftWorx (GE Healthcare) for image acquisition and analyses. Electrical impedance was measured by xCELLigence (OLS) and analyzed by the RTCA (ACEA Biosciences) software. For de novo structure prediction the online PEP-FOLD3 predictor was used. Structural representations were prepared with PyMOL (The PyMOL Molecular Graphics System, Version 1.7.7.6 Schrödinger, LLC).

### Ethics statement

Macrophages were generated from buffy coats of healthy blood donors who give written informed consent for the use of blood-derived products for research purposes (IRB# 507/2017BO1). We do not collect data concerning age, gender or ethnicity and comply with all relevant ethical regulations. All buffy coat donations are received in anonymous form and chosen randomly. Ethics board approval was not required for our work with zebrafish embryos. According to the Directive 2010/63/EU of the European Parliament and of the EU Council on the protection of animals used for scientific purposes, only experiments on freely feeding larvae (i.e. after day 5 of development) are considered “Research on Animals”. Therefore, this definition does not apply to our experiments, which focused on early embryonic development before day 5 of development. In detail, we observed embryonic development until 48 hours post fertilization.

## Author contributions

RBö, RBu, VT, CS, NR, PM and MS designed experiments. RBö, RBu, HP and VT performed experiments. RBö, RBu, HP, NR, PM and MS analyzed the data. DE, KH, JW and MS contributed reagents and analysis tools. MS wrote the manuscript, conceived the overall study and developed the manuscript to its final form. All authors contributed to manuscript editing, read and approved the final manuscript draft.

## Acknowledgements

We are grateful to Christian Sinzger for providing HCMV constructs and giving helpful comments and suggestions. We would further like to thank Ulrich Lauer and his team for support with xCELLigence measurements and the team of the Transfusion Medicine at the University Hospital Tübingen (Taman Bakchoul) for providing buffy coats. This work was in part funded by basic research support given from the University Hospital Tübingen, Medical Faculty as well as an IZKF stipend to Rebecca Böffert.

## Data availability

All data generated and analyzed during this study are included in this published manuscript. Further datasets supporting this study are available from the corresponding author upon request.

